# A method for predicting evolved fold switchers exclusively from their sequences

**DOI:** 10.1101/2020.02.19.956805

**Authors:** Allen K. Kim, Loren L. Looger, Lauren L. Porter

## Abstract

Although most proteins with known structures conform to the longstanding rule-of-thumb that high levels of aligned sequence identity tend to indicate similar folds and functions, an increasing number of exceptions is emerging. In spite of having highly similar sequences, these “evolved fold switchers” (1) can adopt radically different folds with disparate biological functions. Predictive methods for identifying evolved fold switchers are desirable because some of them are associated with disease and/or can perform different functions in cells. Previously, we showed that inconsistencies between predicted and experimentally determined secondary structures can be used to predict fold switching proteins (2). The usefulness of this approach is limited, however, because it requires experimentally determined protein structures, whose magnitude is dwarfed by the number of genomic proteins. Here, we use secondary structure predictions to identify evolved fold switchers from their amino acid sequences alone. To do this, we looked for inconsistencies between the secondary structure predictions of the alternative conformations of evolved fold switchers. We used three different predictors in this study: JPred4, PSIPRED, and SPIDER3. We find that overall inconsistencies are not a significant predictor of evolved fold switchers for any of the three predictors. Inconsistencies between α-helix and β-strand predictions made by JPred4, however, can discriminate between the different conformations of evolved fold switchers with statistical significance (p < 1.7*10^−13^). In light of this observation, we used these inconsistencies as a classifier and found that it could robustly discriminate between evolved fold switchers and evolved non-fold-switchers, as evidenced by a Matthews correlation coefficient of 0.90. These results indicate that inconsistencies between secondary structure predictions can indeed be used to identify evolved fold switchers from their genomic sequences alone. Our findings have implications for genomics, structural biology, and human health.

## Introduction

Relative to the number of well-annotated non-redundant proteins (160 million, (3)), the number of unique protein folds is very small, estimated at ~2000 (4). Because 60-90% of these annotated proteins are estimated to fold into stable three-dimensional structures (5), many different amino acid sequences encoding folded proteins must assume the same fold. This large repertoire of sequences can be envisioned as a map, referred to as “sequence space”, in which proteins with high levels of aligned sequence identity or similarity are close in space, and proteins with dissimilar sequences are farther apart. Studies on large sets of protein structures have shown that proteins even moderately close in sequence space (i.e. with ≥40% aligned identity) tend to adopt the same folds (6).

Recent studies have shown, however, that some proteins very close in sequence space (i.e. with highly similar sequences) can nevertheless adopt different stable folds that sometimes perform different functions (7–9). These proteins, referred to as “evolved fold switchers” (1), can have sequence identities as high as 98% but nevertheless adopt completely disparate folds, such as an α-helical bundle for one protein and a β-barrel for the other (8).

Evolved fold switchers are biologically significant for two reasons. Firstly, these proteins reveal mutational pathways between two protein folds. These pathways support the hypothesis that proteins with novel folds and functions can evolve through stepwise mutation (10, 11). Secondly, because evolved fold switchers challenge the notion that similar amino sequences reliably encode the same protein fold, they could potentially be used to improve methods for protein structure prediction (2) and protein design (12).

To date, three salient examples of evolved fold switchers have been reported in the literature (**Figure 1**). Firstly, the Cro/cI protein superfamily contains two different C-terminal topologies: one α-helix, and the other β-strand (7). This superfamily of transcriptional repressors is present in bacteriophages, and evolutionary analysis suggests that the observed topological differences arose through stepwise mutation (13) as opposed to nonhomologous replacement, a mechanism involving the exchange of a segment of a protein’s ancestral gene for a copy of a nonhomologous gene fragment (13). Although the folds of these proteins appear to be modified through stepwise mutation, the *in vivo* consequences of these changes in secondary and tertiary structure remain uncertain. The second family of evolved fold switchers are the bacterial transcription factors NusG and RfaH (8). These proteins regulate both transcription and translation, albeit by different mechanisms. Both proteins have two domains. The N-terminal domain is a conserved NGN-like domain important for fostering transcriptional readthrough. In NusG, the C-terminal domain (CTD) is a β-barrel involved in Rho-mediated transcription and translation. In contrast, RfaH’s CTD folds into an α-helical bundle that confers additional DNA-binding specificity to the N-terminal domain (8). Remarkably, upon binding both its specific promoter and RNA polymerase, the RfaH CTD refolds into a β-barrel that binds the S10 ribosomal subunit, fostering efficient protein translation (14). The third set of evolved fold switchers, GA98 and GA98, arose from a combination of directed evolution and rational protein design (11, 15). Remarkably, these protein variants are 98% identical (9) but maintain their functions of human serum albumin (HSA) binding (GA98) and IgG binding (GB98).

**Figure 1.**
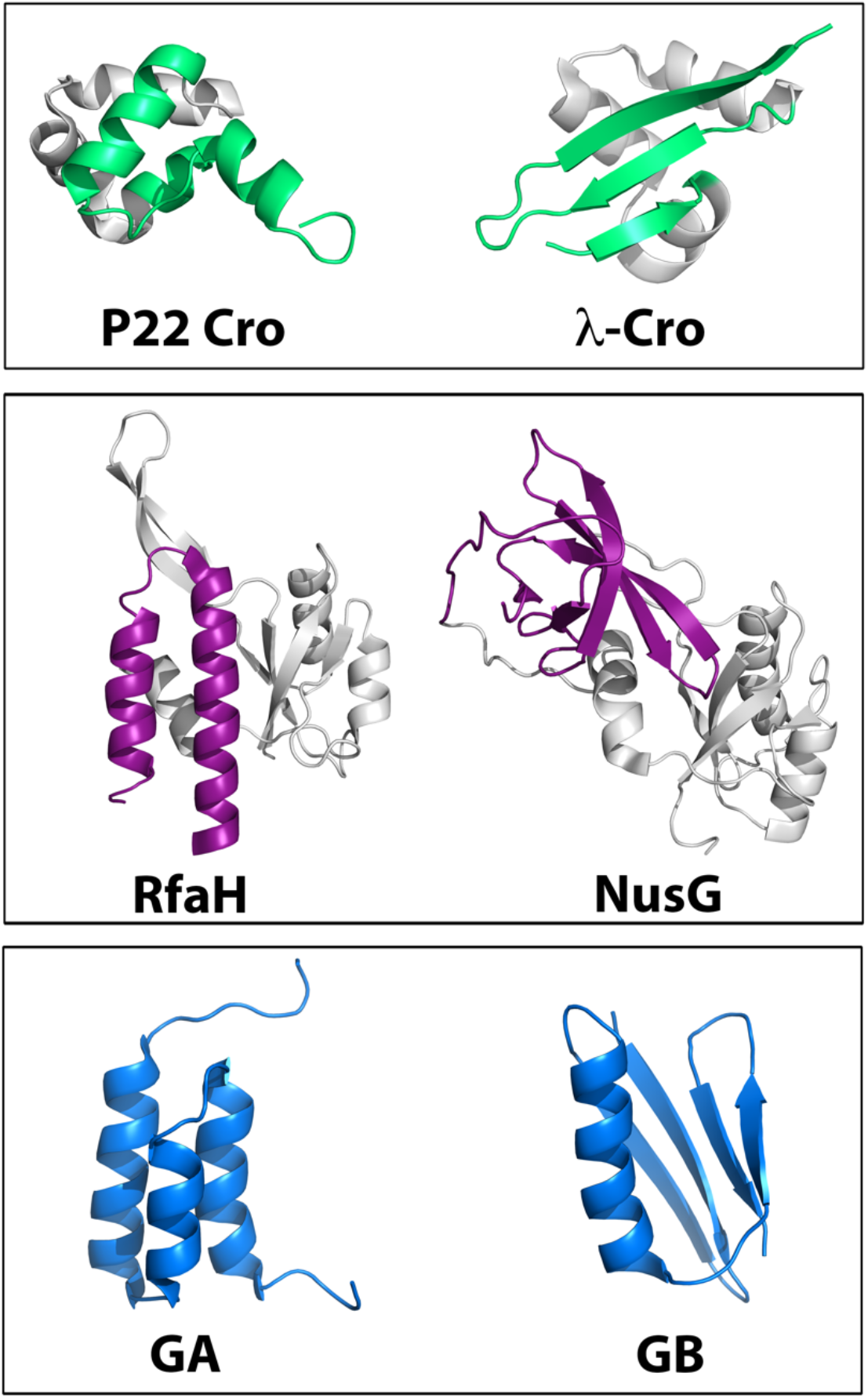
Three sets of evolved fold switchers were used in this analysis: the the p22 and λ-Cro (Cro/cI family) (top), RfaH/NusG family (middle), and engineered proteins GA/GB (bottom). Colored regions of each protein represent regions in which the secondary structures differ between the two proteins whereas gray regions have essentially identical structures. Cartoon diagrams were made with PyMOL.

Predictive methods for identifying more evolved fold switchers are desirable. Although the three current examples provide interesting examples of α<->β transitions accompanied by functional change (at least in the cases of RfaH/NusG and GA/GB), more examples are necessary to understand what kind of mutations tend to foster such radical structural and functional changes. Natural examples of evolved fold switching are especially desirable because of their possible biological relevance: small sequence variations exist between even closely-related eukaryotes, and some evidence suggests that these variations can lead to structural remodeling associated with disease (16) or contribute to cellular behaviors (17). Unfortunately, the current examples of naturally evolved fold switchers were stumbled upon by chance rather than detected by predictive methods. As a result, the rate of discovery has been very slow (3 examples in 60+ years of protein structure determination).

Previously we showed that secondary structure predictions inconsistent with experimentally determined secondary structures can be used to identify extant fold-switching proteins (2). These proteins, in contrast to evolved fold switchers, have a single amino acid sequence that adopts different folds and functions in response to cellular stimuli (1). Thus extant fold switchers have one amino acid sequence that adopts different folds and functions. Inconsistent secondary structure predictions, particularly α<->β inconsistencies, were used to successfully identify extant fold switchers with one solved structure deposited in the PDB. This result suggests that the negative information found in incorrect homology-based secondary structure predictions can be a useful clue for identifying fold switchers. These calculations are severely constrained, however, by the limited number of solved protein structures, which is 3 orders of magnitude smaller than the number of high-quality non-redundant genes currently annotated (3).

Here we test whether inconsistencies between secondary structure predictions can be used to robustly identify evolved fold switchers from their sequences alone. To do this, we used three high-performing homology-based methods to predict the secondary structures of amino acid sequences from the three protein families discussed above. We then looked for inconsistencies between the predictions of the alternatively folded proteins from each family. Upon comparing these inconsistencies with those calculated from three protein families with no evolved fold switching observed (termed single-fold proteins), we found that inconsistent secondary structure predictions alone were not sufficient to identify evolved fold switchers from their sequences. In contrast, α/β prediction discrepancies from JPred4—but not the other two algorithms—appeared to discriminate between fold switchers and single-fold proteins with a high level of statistical significance. We used these discrepancies to develop a classifier and determined that it had a Matthews Correlation Coefficient of 0.90, suggesting that α/β prediction discrepancies from JPred4 could potentially identify evolved fold switchers from their genomic sequences with high confidence. Thus, this approach could be used to search protein sequence space for more evolved fold switchers.

## Results

### Secondary structure discrepancies are not good predictors of evolved fold switchers

First, we tested whether secondary structure predictors could identify appreciable differences between the distinct folds of evolved fold switchers. To do this, we ran three secondary structure prediction algorithms—JPred4 (18), PSIPRED (19), and SPIDER3 (20)—on all solved structures of RfaH/NusG, Cro/cI proteins, and designed GA/GB variants. **Figure 2** shows examples of discrepancies found for each of the three protein families. In all three cases, significant differences between predictions of the different conformations are evident. Most notably, JPred4 correctly predicted single mutations in the GA/GB system that lead to drastic changes in secondary structure. We note that these secondary structure predictions are not particularly consistent with the solved protein structures of any of the evolved fold switchers. Previous work has shown the disagreement between predicted and experimentally determined secondary structure is a possible approach for identifying extant fold switchers (2), but here we focus on the differences between the predictions themselves because this approach potentially allows for evolved fold switchers to be identified from their genomic sequences alone, without the need for a previously solved protein structure.

**Figure 2.**
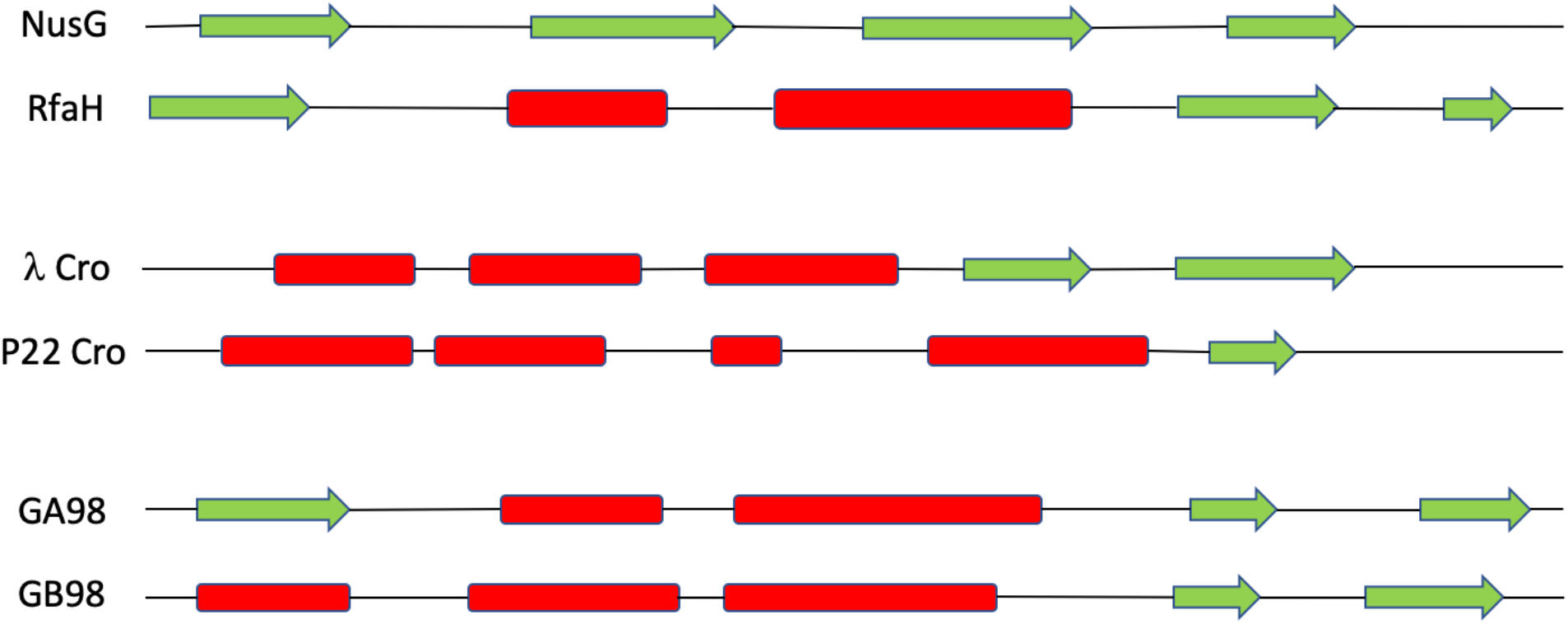
Secondary structure predictors identify secondary structure discrepancies in all three groups of evolved fold switchers. Green arrows represent β-strands, red boxes represent α-helices, and black lines represent coils. All predictions were made by JPred4 on the following PDBs: NusG—2LQ8 (last 50 residues); RfaH—2OUG (last 50 residues); λ-Cro (1COP, full sequence), p22 Cro (1RZS, full sequence), GA98 (2LHC, full sequence), GB98 (2LHD, full sequence).

We then quantified these differences and found that most of them gave the typical level of error for homology-based secondary structure predictors (~20%, (21)) (**Table 1**). To do this, we calculated the average Q3 value between all proteins from the same family with different experimentally determined folds. The Q3 value gives a binary score of 1/0 for agreement/disagreement between the predicted and experimentally determined secondary structure of each amino acid position (<u>H</u>elix, <u>E</u>xtended β-strand, or <u>C</u>oil); this score is summed over sequence and normalized by length. **Table 1** shows that among the nine discrepancies calculated, six were within error of the expected discrepancy level of 20% or less, meaning a Q3 accuracy of 80% or more, the typical value for most homology-based secondary structure predictors (21). The three exceptions were JPred’s predictions of the RfaH/NusG family (45±0.81%) and p22/λ family (30±3.0%) as well as SPIDER3’s predictions of the p22/λ family (27±4.2%).

**Table 1.** Overall prediction discrepancies (mean ± median) for fold switchers

|  | RfaH/NusG | Cro/cI | GA/GB |
| --- | --- | --- | --- |
| JPred4 | $45 \pm 0.81\%$ | $31 \pm 2.1\%$ | $26 \pm 7.5\%$ |
| PSIPRED | $17 \pm 2.2\%$ | $23 \pm 4.4\%$ | $19 \pm 6.1\%$ |
| SPIDER3 | $18 \pm 0.81\%$ | $27 \pm 4.2\%$ | $20 \pm 6.8\%$ |

Because two-thirds of the calculated discrepancies between evolved fold switchers were within the error expected for homology-based secondary structure predictions of single-fold proteins, we hypothesized that secondary structure predictions of single-fold proteins would yield similar levels of discrepancy. To test this, we ran all three secondary structure predictors on sequences from experimentally determined structures of proteins with very different folds: helical bundles, ubiquitin-like folds, and SH3-like domains. **Table 2** shows that, indeed, the overall prediction discrepancies for these three single-fold protein families were largely similar to those of evolved fold switchers; almost all predictions were within the 20% level of expected error, though the largest level of discrepancy came out slightly above that (29±4.6% for SPIDER3 calculations of the ubiquitin-like fold). These results indeed suggest that secondary structure discrepancies among sequences of related proteins are not a good predictor of fold switching.

**Table 2.** Overall prediction discrepancies (mean ± median) for single-fold proteins

|  | SH3 | Helical bundle | Ubiquitin-like |
| --- | --- | --- | --- |
| JPred4 | 13±6.0% | 25±6.6% | 25±2.6% |
| PSIPRED | 14±6.9% | 23±5.7% | 24±5.2% |
| SPIDER3 | 20±5.3% | 25±4.3% | 29±4.6% |

### Secondary structure prediction discrepancies can identify evolved fold switchers

Although overall secondary structure prediction discrepancies do not appear to reveal evolved fold switchers, we sought to determine whether α<->β discrepancies might be more predictive. Previous work suggests that these discrepancies can be enriched in fold switchers compared to single-fold proteins (2). To do this, we quantified the number of α<->β discrepancies between evolved fold switchers and normalized them by the minimum number of secondary structure annotations from one of the two folds. **Table 3** shows that JPred4 yields significantly higher α<->β discrepancies than the previously published value of 6% for single-fold proteins (2). The other two predictors gave much lower frequencies of α<->β discrepancies, mostly within error of 0%, with the notable exception of SPIDER3’s Cro:cI value of 15±4.8%.

**Table 3.**
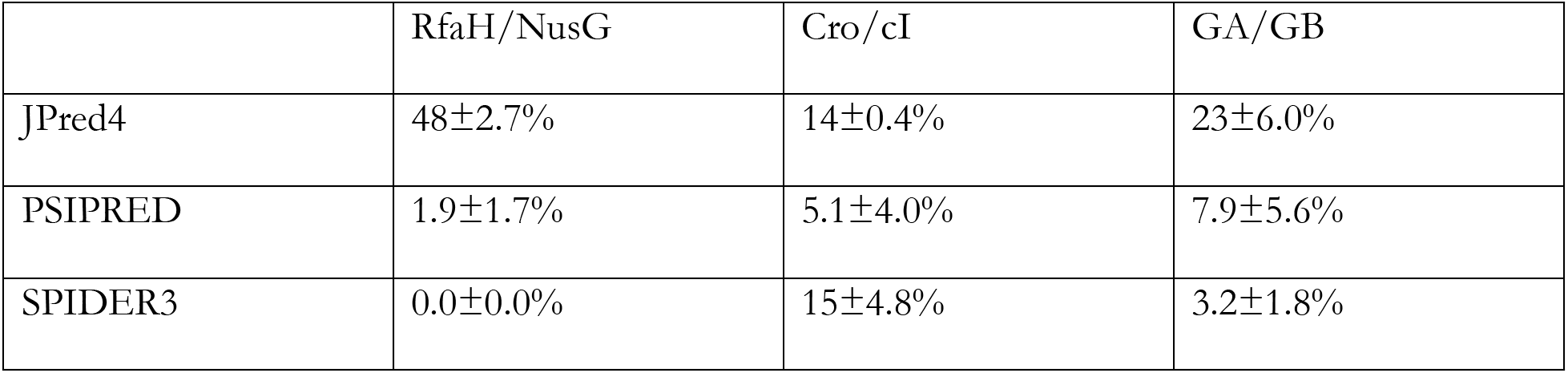
α<->β prediction discrepancies (mean ± median) for fold switchers

|  | RfaH/NusG | Cro/cI | GA/GB |
| --- | --- | --- | --- |
| JPred4 | 48±2.7% | 14±0.4% | 23±6.0% |
| PSIPRED | 1.9±1.7% | 5.1±4.0% | 7.9±5.6% |
| SPIDER3 | 0.0±0.0% | 15±4.8% | 3.2±1.8% |

To test whether JPred4 systematically yields higher levels of α<->β discrepancies, and to test the performance of the other two secondary structure predictors, we reran these calculations on our three selected families of single-fold proteins. **Table 4** shows that JPred4 yields significantly lower α<->β discrepancies for single-fold proteins than for evolved fold switchers, suggesting that it may have the potential to identify evolved fold-switching proteins from their genomic sequences. In contrast, α<->β prediction discrepancies between evolved fold switchers and single-fold proteins were largely within error of zero for both PSIPRED and SPIDER3.

**Table 4.** α<->β prediction discrepancies (mean ± median) for single-fold proteins

|  | SH3 | Helix | Ubiquitin-like |
| --- | --- | --- | --- |
| JPred4 | 0.29±0.80% | 7.9±4.3% | 5.0±1.9% |
| PSIPRED | 2.2±3.1% | 0.99±1.1% | 5.0±5.3% |
| SPIDER3 | 2.2±2.6% | 4.6±4.2% | 10±3.7% |

### JPred4 as a potential classifier for fold switching proteins

Because JPred4 seems to discriminate between evolved fold switchers and evolved single-fold proteins, we tested its robustness as a classifier. To do this, we compared the distributions of α/β prediction discrepancies for both evolved fold switchers and evolved single-fold proteins (**Figure 3**). We found that the distributions were very different, with p < 1.7*10^−13^ (Kolmogorov-Smirnov test). Over 95% of α<->β prediction discrepancies for single-fold proteins had values of less than 10%. In contrast, <5% of α<->β prediction discrepancies fell under the 10% threshold. In light of this observation, we calculated the Matthews Correlation Coefficient (22) for fold switchers and single-fold proteins at a 10% threshold and found it to be 0.90, in good agreement with the experimental data. This result suggests that α<->β prediction discrepancies calculated from JPred4 could indeed be a good predictor for evolved fold switchers.

**Figure 3.**
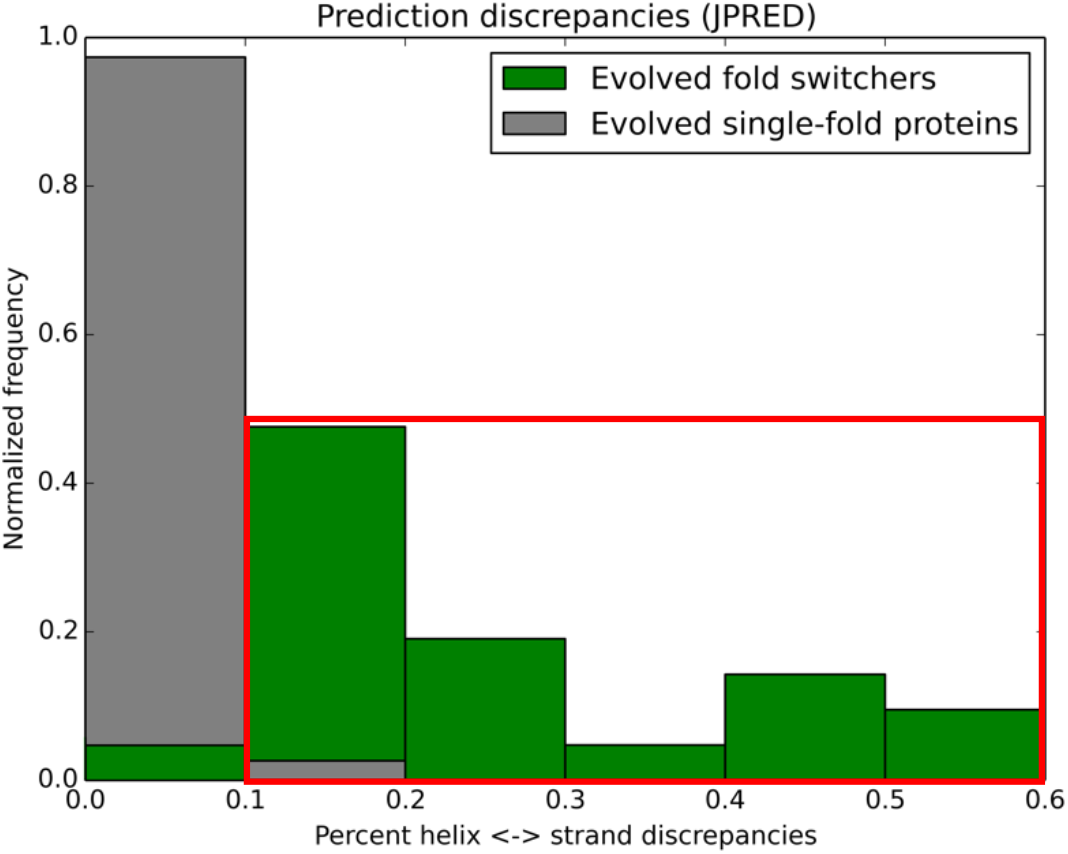
Evolved fold switchers have significantly higher proportions of α<->β discrepancies than evolved single-fold proteins. These distributions differ with a probability < 1.7*10^−13^ by the Kolmogorov-Smirnov test. Levels of discrepancy exceeding 10% are boxed in red. Using the 10% line as a threshold for a classifier, α<->β discrepancies yield a Matthew’s Correlation Coefficient of 0.90.

## Discussion

Although most proteins with high levels of sequence identity tend to adopt the same folds and perform the same functions, a number of exceptions, called evolved fold switchers (1), have emerged. Accurately predicting evolved fold switchers from their genomic sequences would be useful because there are biologically relevant examples in which one or several mutations in a protein can lead to structural changes associated with disease (16) or different cellular behaviors (17). We also hypothesized that this approach could be more successful on evolved fold-switchers than on extant fold switchers because evolved fold switchers tend to have only one visible folded state by NMR (9, 14, 23, 24), and they may only adopt a single fold, as in the cases of the Cro/cI proteins and some GA/GB variants. Proteins with more than one detectable ground state, such as lymphotactin (25), might be more difficult to predict with this method because it lacks a highly stable fold different from its homologs. This structural ambivalence could confound homology-based predictors by obscuring appreciable changes in secondary structure.

We note that although α<->β discrepancies appear to be a good predictor for evolved fold switchers, overall structural discrepancies could not distinguish evolved fold switchers from single-fold protein families. These results are consistent with previous work showing that secondary structure predictions tend to predict coil least accurately. These results are also largely consistent with our previous work (2), which showed that coil <-> helix or coil <-> strand discrepancies were more common for non-fold-switching (*i.e.* single-fold) regions of proteins. One other area of consistency is that extant fold switchers tend to be enriched in α<->β discrepancies. One surprising observation, however, is that α<->β discrepancies were enriched for extant fold switchers among three secondary structure predictors. Extant fold switchers—unlike some evolved fold switchers--have single sequences that can remodel their secondary structures in response to cellular stimuli. Here, only JPred4 appears to produce significant α<->β discrepancies for extant fold switchers. It is not immediately obvious why Jpred4 appears to have stronger predictive power than PSIPRED or SPIDER3 in this case. All three are neural networks that derive their results from position-specific scoring matrices produced by PSI-BLAST (26). One possibility could be that the three algorithms were trained on different sets of proteins.

While this method shows promise for identifying evolved fold switchers from their genomic sequences, further work is needed to validate the approach. Determining the structures of related sequences with discordant secondary structures would be best. Nevertheless, this method does present advantages over our previous work because it does not require any solved protein structures to make predictions. It does require a number of related sequences, however. For that reason, this method could not be used on orphan proteins (27), for example. We are also unsure of its ability to identify extant fold switchers, though it does robustly identify RfaH, which can be described as both an extant fold switcher because it can remodel its secondary structure and an evolved fold switcher because its α-helical conformation appears dominant by NMR, but it has homologs that fold into β-barrels. Regardless, we anticipate that the additional information gained through identification of more fold switchers—identified by this method or others—will inform more robust and generalizable predictions in the future.

## Methods

### Secondary structure predictions of evolved fold switchers

All amino acid sequences from 21 evolved fold switchers were downloaded from the Protein Data Bank (PDB) and saved as individual FASTA (28) files. Separate secondary structure predictions were run on each file using JPred4, PSIPRED, and SPIDER3. JPred4 predictions were run remotely using a publicly downloadable scheduler available on the JPred4 website (compbio.dundee.ac.uk/jpred/). SPIDER3 calculations were performed on the Spark’s Lab webserver (https://sparks-lab.org/server/spider3). PSIPRED calculations were run locally using the Uniprot90 database ((29), 11/12/2019). Secondary structure predictions from .jnetpred (JPred), .horiz files (PSIPRED), and .spd33 files (SPIDER3) were converted into FASTA format. Each residue was assigned one of three secondary structures: “H” for helix, “E” for extended β-strand, and “C” for coil. Chain breaks were annotated “-.” PDB IDs from each family of evolved fold switchers were as follows: Cro/cI: 1RZS, 2HIN, 3BD1, 2OVG, 5W8Y, 2PIJ, 5W8Z, 1COP; NusG/RfaH: the last 50 residues of 2OUG, 1NPP, 1NZ9, 2LQ8, 1MI6; GA/GB: 2LHC, 2LHG, 2KDL, 2JWS, 2JWU, 2KDM, 2LHD, 2LHE. Cartoon diagrams of these evolved fold switchers (**Fig. 1**) were made using Pymol (30).

### Secondary structure prediction accuracy calculations

To calculate secondary structure accuracies, we used the 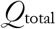 (or 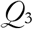) metric (31), which compares predicted and experimentally determined secondary structures one-by-one, residue-by-residue. Predictions were scored as follows: (in)consistent pairwise predictions are assigned a score of (0)/1, summed, and normalized by the minimum length of the two sequences compared. We did not include chain breaks for either scoring or normalization. Although 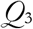 values are typically expressed as decimals, here we report them as percentages. Registers of secondary structure predictions were determined by multiple sequence alignments of protein families calculated by ClustalOmega (32).

### Single-fold protein families

We obtained our single-fold protein families from the CATH database (33), release 1/11/20. Specifically, we used domains with <35% identity from 3.10.20 (ubiquitin-like roll), 1.10.8 (helicase, helical bundle), and 2.30.30.40 (SH3 domains).

### Helix <-> strand discrepancies and distribution

Helix <-> strand discrepancies were normalized by the minimum number of secondary structure annotations (helix or sheet) in the two sequences compared. The distributions in Figure 3 were plotted using Matplotlib (34).

